# The “ForensOMICS” approach to forensic post-mortem interval estimation: combining metabolomics, lipidomics and proteomics for the analysis of human bone

**DOI:** 10.1101/2022.09.29.510059

**Authors:** Andrea Bonicelli, Hayley L. Mickleburgh, Alberto Chighine, Emanuela Locci, Daniel J. Wescott, Noemi Procopio

**Affiliations:** School of Natural Sciences, University of Central Lancashire, Preston, PR1 2HE, United Kingdom; The Forensic Science Unit, Faculty of Health and Life Sciences, Ellison Building, Northumbria University, Northumbria University Newcastle, Newcastle Upon Tyne, NE1 8ST, United Kingdom; ACASA – Department of Archaeology, Faculty of Humanities, University of Amsterdam, PO Box 94203, 1090 GE, Amsterdam, The Netherlands; Forensic Anthropology Center, Texas State University, San Marcos 78666, Texas, United States; Department of Medical Science and Public Health, Section of Legal Medicine, University of Cagliari, Monserrato, 09042, Italy

**Keywords:** human bone, post-mortem interval, decomposition, multi omics, metabolomics, lipidomics, proteomics

## Abstract

The combined use of multiple omics methods to answer complex system biology questions is growing in biological and medical sciences, as the importance of studying interrelated biological processes in their entirety is increasingly recognized. We applied a combination of metabolomics, lipidomics and proteomics to human bone to investigate the potential of this multi-omics approach to estimate the time elapsed since death (i.e., the post-mortem interval, PMI). This “ForensOMICS” approach has the potential to improve accuracy and precision of PMI estimation of skeletonized human remains, thereby helping forensic investigators to establish the timeline of events surrounding death. Anterior midshaft tibial bone was collected from four female body donors in a fresh stage of decomposition before placement of the bodies to decompose outdoors at the human taphonomy facility managed by the Forensic Anthropological Center at Texas State (FACTS). Bone samples were again collected at selected PMIs (219, 790, 834 and 872 days). Liquid chromatography mass spectrometry (LC-MS) was used to obtain untargeted metabolomic, lipidomic and proteomic profiles from the pre- and post-placement bone samples. Multivariate analysis was used to investigate the three omics blocks by means of Data Integration Analysis for Biomarker discovery using Latent variable approaches for Omics studies (DIABLO), to identify the reduced number of markers that could effectively describe post-mortem changes and classify the individuals based on their PMI. The resulting model showed that pre-placement bone metabolome, lipidome and proteome profiles were clearly distinguishable from post-placement profiles. Metabolites associated with the pre-placement samples, suggested an extinction of the energetic metabolism and a switch towards another source of fuelling (e.g., structural proteins). We were able to identify certain biomolecules from the three groups that show excellent potential for estimation of the PMI, predominantly the biomolecules from the metabolomics block. Our findings suggest that, by targeting a combination of compounds with different post-mortem stability, in future studies we could be able to estimate both short PMIs, by using metabolites and lipids, and longer PMIs, by including more stable proteins.

## 1 Introduction

The modifications that occur to the human body after death are complex and known to be affected by a variety of intrinsic and extrinsic factors. The rate of decomposition can vary significantly depending on the environment and even the manner of death. Nonetheless, the process of decomposition has been demonstrated to be predictable, providing opportunities to estimate the time elapsed since death (also known as post-mortem interval, PMI) based on gross morphological and/or microscopic changes to the body. Precise and accurate estimation of the PMI is crucial to help establish the timeline of events surrounding death and can help medicolegal investigators with the identification of the deceased and can corroborate or negate other forensic evidence.

In the first hours after death, the body undergoes several post-mortem changes, including progressive cooling (*algor mortis*), increased rigidity associated with muscle stiffness (*rigor mortis*), and pink-purplish discolouration, in light skinned individuals, caused by the lack of blood circulation in and settling of blood in the lowest areas (*livor mortis*)^1–3^. After these stages, as the time since death increases, the breaking down and liquefaction of the organs and other soft tissues will occur: a process referred to as putrefaction. The lack of oxygenated circulation induces cellular hypoxia, leading to swelling of the cells, and subsequent rupture of cell membranes and releasing of digestive enzymes. This triggers autolytic digestion of the soft tissues^4^. The body becomes fully anaerobic, allowing anoxic (endogenous) bacteria to proliferate and transmigrate throughout the entire body^5,6^. The activity of endogenous bacteria results in the accumulation of gases which cause bloating of the soft tissues, starting in the abdomen, but al also visible in the face in early decomposition stages, and progressing towards the rest of the body. Colonisation of the body by insects and exogenous bacteria, mostly aerobic microorganisms, contributes further to the changes and reduction of the soft tissues^7,8^. Besides these, other extrinsic factors including abiotic environmental conditions (*e*.*g*., humidity, temperature, sun exposition, aeration, burial context) and biotic factors, such as the presence and type of microorganisms, insects, and scavengers^9,10^, will affect the rate of decomposition of the soft tissues. Intrinsic factors known to affect the rate of decomposition include, among others, body mass index, and antemortem and perimortem pathological conditions^11^. Completion of putrefaction and the activity of insects consuming the decomposing soft tissues, will leave the remains completely, or almost completely, skeletonized, and dry.

The complex nature and interplay of intrinsic and extrinsic variables involved in the process of decomposition, means that developing accurate and precise models for PMI estimation is extremely challenging. Traditional methods of PMI estimation include calculating PMI using the body temperature and ambient temperature (which relies on the predictability of *algor mortis*, and works for short PMIs only), or the visual assessment of gross morphological changes to the body to estimate a PMI range (short and longer PMIs). Since the rate of gross morphological changes is variable, methods that rely on visual scoring of decomposition stages suffer from issues of poor accuracy and precision. An additional problem of such methods is the effect of interobserver variation on the scoring of decomposition stages. For all commonly used PMI estimation methods, the accuracy and precision decreases considerably as decomposition progresses, and is particularly problematic when the remains are partially or completely skeletonized^2,3^.

In recent years, the number of studies exploring the use of biomolecular methods of PMI estimation has risen sharply, due to their potential for providing more accurate and precise estimation methods based on the rates of decay of different molecules and compounds^12–16^. Better understanding of biomolecular decomposition of bone will provide opportunities to develop biomolecular methods for estimation of longer PMIs (*i*.*e*., timeframes in which soft tissues are unlikely to be preserved). Moreover, through the combined analysis of multiple different panels of omics, greater precision and accuracy of PMI estimation can potentially be achieved.

Biomolecular decomposition is caused by both enzymatic and microbial breakdown of large molecules, resulting in the breakage of proteins into amino acids (AA), of carbohydrates into more simple monosaccharides, and of lipids into simpler fatty acids chains^17,18^. In carbohydrate decomposition, the complex polysaccharides are normally broken down via microbial activity into smaller units of monosaccharides. This breakdown can be achieved by oxidation that produces carbon dioxide and water and can partially decompose resulting in the production of organic acids and alcohols. Alternatively, the monosaccharides can be degraded by fungal activity into glucuronic, citric, and oxalic acids, or by bacteria into lactic, butyric, and acetic acids^17,19^. During decay of lipids, free saturated and unsaturated fatty acids are released due to hydrolysis mediated by the action of intrinsic lipases released after death. These can then be converted into hydroxyl fatty acids (the main constituent of adipocere) by the action of specific bacterial enzymes in humid environments, or can associate with potassium and sodium ions, resulting in the formation of salts^19^. Protein degradation is primarily an enzyme-driven process, led by the action of proteases, which occurs at different rates for different proteins and tissues. Proteolytic enzymes induce the hydrolytic breakdown of proteins and the production respectively of proteoses, peptones, polypeptides, and finally AA, which can be further modified via deamination (production of ammonia), decarboxylation (production of cadaverine, putrescine, tyramine, tryptamine, indole, skatole and carbon dioxide) and desulfhydralation (production of hydro gen sulphide, pyruvic acid, and thiols)^17,19^.

The analysis of low molecular weight compounds and decomposition by-products is becoming more popular in forensic science, particularly for the purpose of estimating the PMI^20^. Time since death was recently reported as the main variable driving modifications in the metabolome occurring after death^21^ in many soft tissues and fluids, so the metabolomic appears ideal to estimate PMI. However, the potential forensic significance of the post-mortem bone metabolome is as yet underexplored^22^. Several studies on soft tissues (vitreous and aqueous humour) have examined metabolomics for the purpose of determining short PMIs. Examining longer PMIs based on metabolomics analysis of humour has not been possible due to evaporation and leakage through the corneal surface as time since death progresses^15^. Girela et al.^23^ reported a significant positive correlation between post-mortem interval and taurine, glutamate, and aspartate levels observed in vitreous humour. These results were partially confirmed by Zelentsova et al.^16^, who found a correlation between the levels of hypoxanthine, choline, creatine, betaine, glutamate, and glycine and PMI. Another approach employing ^1^H-NMR on aqueous humour from pig heads reported taurine, choline, and succinate as major metabolites involved in the post-mortem modification^15^. The same study also showed an orthogonally constrained PLS2 (oCPLS2) model showing prediction error of 59 min for PMI < 500 min, 104 min for PMI from 500 to 1000 min, and 118 min for PMI > 1000 min. Beside humour, muscle is one of the most frequently targeted tissues in metabolomics studies focused on short PMI estimation. Pesko et al.^14^ recently evaluated rat and human biceps femoris muscles from the same individuals at different PMIs, demonstrating an increase of the abundance of several metabolites, including most of those derived from the breakdown of proteins, and in particular highlighting how threonine, tyrosine, and lysine show the most consistent and predictable variations in relatively short PMIs. An untargeted metabolomics study on muscle tissue also indicated the potential of isolating biomarkers associated with age^24^, suggesting the potential applications of metabolomics for both age-at-death (AAD) and PMI estimation. To date, only three studies have used lipidomics assays for PMI estimation. Two of them were conducted on muscle tissue and showed, in general, a negative correlation between most lipid classes and PMI, as well as an increment in free fatty acids^25,26^. The third study applied lipidomics to trabecular bone samples from calcanei spanning a PMI of approximately seven years and highlighted the presence of 76 potential N-acyl AA that could be employed for PMI estimation, however their correlation with PMI has not yet been fully elucidated^27^.

Several studies have tried to quantify the degree of survival of proteins and the accumulation of post-translational modifications (PTMs) of AA in both animal and human models^10,11,13,28,29^ as well as under different conditions (*e*.*g*., in aquatic environments, different types of coffins, buried vs. surface)^28–30^. The premise of these studies is that the protective action of the hydroxyapatite is expected to enhance the survival of proteins, allowing potential estimation of longer PMIs. Results generally showed that blood/plasma and ubiquitous proteins decrease in their abundance constantly starting from the early decomposition stages, whereas proteins more strongly connected to the mineral matrix such as bone-specific proteins are able to survive for longer PMIs and can be useful indicators for PMI estimation also in skeletonised remains. Similarly, also the accumulation of specific non-enzymatic PTMs, such as deamidations, can be used as a biomarker for the evaluation of the PMI in bones.

While many studies have applied different analytical platforms for proteomics, metabolomics and lipidomics to several different matrices^14–16,23,31–35^, relatively little is known about the biomolecular decomposition of bone tissue. Moreover, while clinical studies have applied multi-omics methods with some frequency, their potential for development of more precise and accurate biomolecular PMI estimation methods is has not been explored. The present study applies, for the first time, a multi-omics approach (*i*.*e*., combined proteomics, metabolomics and lipidomics, defined here as the “ForensOMICS” approach) to pre- and post-decomposition tibial cortical bone samples from four human female body donors, to identify potential multi-omics biomarkers of time since death. The multi-omics approach uses the natural differences in manner and rate of decomposition between the different biomolecules (proteins, metabolites, lipids) to expand the potential range of PMIs and to cross-correlate results between different sets of biomarkers to narrow down PMI ranges based on the degradation of multiple biomolecules. The use a of a single omics technique would not be suitable to investigate a wide range of potential PMIs.

Metabolites and lipids are appropriate for short PMIs while protein have been proved to be stable across longer ones. Therefore, the combination of the three classes of biomolecules aims to obtain ideal coverage across a wider range of PMIs. Additional advantages of the combined application of these methods potentially include greater flexibility in application across different environments and different post-mortem treatments since the use of multiple types of biomolecules and compounds increases the likelihood of retrieving suitable markers for PMI estimation. The present study provides proof-of-concept for future validation of the multi-omics approach on a larger number of individuals.

## 2 Materials and Methods

### 2.1 Body Donors

Bone samples were collected from four female human body donors, aged between 61 and 91 years (mean 74±11.6 SD), at the Forensic Anthropology Center at Texas State University (FACTS). FACTS receives whole body donations for scientific research under the Texas revised Uniform Anatomical Gift Act^36^. Body donations are made directly to FACTS and are exclusively acquired through the expressed and documented will of the donors and/or their legal next of kin. Demographic, health, and other information are obtained through a questionnaire completed by the donor or next of kin. The data are securely curated by FACTS, and the body donation program complies with all legal and ethical standards associated with the use of human remains for scientific research in the United States. The number of individuals (n=4) used in this preliminary study is consistent with other taphonomic studies conducted on human remains for proof-of-concept purposes. Larger sample sizes may be used to validate preliminary results, such as those proposed by this study, at a later stage.

The bodies were stored in a cooler at 4°C prior to sampling. After collection of the initial (pre-placement) bone samples, the bodies were placed outdoors to decompose at the Forensic Anthropology Research Facility (FARF), the human taphonomy facility managed by FACTS, between April 2015 and March 2018. Two of the four body donors (D1 and D4, see **Tab. 1**), were placed in shallow hand-dug pits which were left open throughout the duration of the decomposition experiment. The pits were covered with metal cages to prevent disturbance by large scavengers. Donors D2 and D3 were deposited in similarly sized hand-dug pits and were immediately buried with soil.

### 2.2 Sampling

Bone samples (ca. 1 cm^3^) of the anterior midshaft tibia (left) were collected prior to placement of the body outdoors, and again upon retrieval of the completely skeletonized remains (right). Each body was in “fresh” stage of decomposition when pre-placement samples were taken, and in “skeletonization” stage when post-placement samples were collected, based on scoring of the gross morphological changes^37^. The duration of each placement and the deposition context are reported in **Table 1**. The soft tissue was incised with a disposable scalpel, and a 12 V Dremel cordless lithium-ion drill with a diamond wheel drill bit was used at max. 5000 revolutions to collect ∼1 cm^3^ of bone. Sampling instruments were cleaned with bleach and deionised water between each individual sample collection. A total of eight samples were collected in Ziploc bags, transferred immediately to a −80°C freezer, and subsequently shipped overnight on dry ice to the Forensic Science Unit at Northumbria University, U.K. The samples were then transferred to a lockable freezer at − 20°C as per UK Human Tissue Act regulations (licence number 12495). Part of the analyses were conducted by the “ForensOMICS” team (N.P. and A.B.) at Northumbria University prior to their transfer to the University of Central Lancashire. Specifically, the bone samples were defrosted, and fine powder was obtained with a Dremel drill equipped with diamond-tipped drill bits operated at speed 5000 rpms, to avoid heat damage caused by the friction with the bone. The collected powder was homogenised and stored in 2 mL protein LoBind tubes (Eppendorf UK Limited, Stevenage, UK) at −80°C until extraction and testing. The powder sample was later divided into 25 mg aliquots. Three biological replicates (*e*.*g*., three aliquots of bone sample per specimen) were extracted and analysed for each specimen. The research and bone sample analyses were reviewed and approved by the Ethics committee at Northumbria University (ref. 11623).

**Table 1.** Sample composition, demographics, deposition context, and PMI. The Sample ID column reports the biological replicates used. Additional information on the body donors and observations made during collection of bone samples (e.g., medical treatments, bone colour and density) can be found in the supplementary information in Mickleburgh et al.^11^.

| Sample ID | Sex | Age (years) | PMI | Deposition context |
| --- | --- | --- | --- | --- |
| Pre-Burial samples |  |  |  |  |
| D1_TF_A | Female | 91 | 10 days | Open pit |
| D1_TF_B | Female | 91 | 10 days | Open pit |
| D1_TF_C | Female | 91 | 10 days | Open pit |
| D2_TF_A | Female | 67 | 2 days | Burial |
| D2_TF_B | Female | 67 | 2 days | Burial |
| D2_TF_C | Female | 67 | 2 days | Burial |
| D3_TF_A | Female | 61 | 3 days | Burial |
| D3_TF_B | Female | 61 | 3 days | Burial |
| D3_TF_C | Female | 61 | 3 days | Burial |
| D4_TF_A | Female | 77 | 10 days | Open pit |
| D4_TF_B | Female | 77 | 10 days | Open pit |
| D4_TF_C | Female | 77 | 10 days | Open pit |
| Post-burial samples |  |  |  |  |
| D1_TS_A | Female | 91 | 219 days | Open pit |
| D1_TS_B | Female | 91 | 219 days | Open pit |
| D1_TS_C | Female | 91 | 219 days | Open pit |
| D2_TS_A | Female | 67 | 790 days | Burial |
| D2_TS_B | Female | 67 | 790 days | Burial |
| D2_TS_C | Female | 67 | 790 days | Burial |
| D3_TS_A | Female | 61 | 834 days | Burial |
| D3_TS_B | Female | 61 | 834 days | Burial |
| D3_TS_C | Female | 61 | 834 days | Burial |
| D4_TS_A | Female | 77 | 872 days | Open pit |
| D4_TS_B | Female | 77 | 872 days | Open pit |
| D4_TS_C | Female | 77 | 872 days | Open pit |

### 2.3 Biphasic extraction, adapted Folch protocol

Chloroform (Chl), AnalaR NORMAPUR^®^ ACS was purchased from VWR Chemicals (Lutterworth, UK). Water Optima™ LC/MS Grade, Methanol (MeOH) Optima™ LC/MS Grade, Pierce™ Acetonitrile (ACN), LC-MS Grade and Isopropanol (IPA), Optima™LC/MS Grade were purchased from Thermo Scientific (Hemel Hempstead, United Kingdom). In total three biological replicates for each of the eight specimens were extracted according to a modified Folch et al.^38^ as follow: 25 mg of bone powder was placed in tube A and 750μL of 2:1 (v/v) Chl:MeOH were added, vortexed for 30s and sonicated in ice for additional 20 min. 300μL of LC-MS grade water was added to induce phase separation and sonicate for another 15 mins. The sample were then centrifuged at 10°C for 5 mins at 2000 RPM. The respective upper and lower fractions were collected and transferred to fresh Eppendorf tubes and the samples were re-extracted with a second time using 750μL of 2:1 (v/v) Chl:MeOH. The two respective fractions were combined and concentrated. The organic lipid fraction was preconcentrated using a vacuum concentrator at 55oC for 2.5 hours or until all organic solvents has been removed. The aqueous metabolite fractions were flash frozen in liquid nitrogen and preconcentrated using a lyophilizer cold trap −65°C over night to remove all water content. The respective dry fractions were then stored at −80 until analysis. The metabolite fraction was resuspended in 100μL in 95:5 ACN/water (v/v) and sonicated for 15 mins and centrifuged for 15 min at 15K RPM at 4°C and supernatant was then transferred to 1.5mL autosampler vials with 200μL microinsert and caped. 20μL of each sample were collected and pooled to create the pooled QC. The lipid extracts were resuspended in 100μL of 1:1:2 (v/v) water:ACN:IPA and sonicated for sonicated for 15 min and centrifuged for 15 min at 15K RPM at 10oC and supernatant was then transferred to 1.5mL autosampler vials with 200μL microinsert and caped. 20μL of each sample were collected and pooled to create the pooled QC. The sample set was then submitted for analysis.

### 2.4 LC-MS analysis

Metabolite and lipid characterization of the bone samples was performed on a Thermo Scientific (Hemel Hempstead, United Kingdom) Vanquish Liquid Chromatography (LC) Front end connected to IDX High Resolution Mass Spectrometer (MS) system. Full details for both metabolomics and lipidomics runs are reported below.

#### 2.4.1 Metabolomics

Hydrophilic Liquid Interaction Chromatography (HILIC) was used for the chromatographic separation for metabolites. The separation was achieved using a Waters Acquity UPLC BEH amide column (2.1 × 150mm with particle size of 1.7μm, part no. 186004802), operating at 45°C with a flow rate of 200μL/min. The LC gradient consists of a binary buffer system, namely buffer “A” (LC/MS grade water) and buffer “B” (LC/MS grade ACN) both containing 10 mM ammonium formate. Independent buffer systems were used for positive and negative electrospray ionisation (ESI) acquisition respectively, for ESI+ the pH of buffers was adjusted using 0.1% formic acid and for negative using 0.1% ammonia solution. The LC gradient was the same for both polarities, namely 95% “B” at T0 hold for 1.5 min and a linear decrease to 50% “B” at 11 min, followed by hold for 4 mins, return to starting condition and hold for further 4.5 mins (column stabilization). The voltage applied for ESI+ and ESI-was 3.5 kV and 2.5 kV respectively. Injection volumes used were 5 μL for ESI+ and 10 μL for ESI-.

#### 2.4.2 Lipidomics

Standard reverse phase chromatography was used for the chromatographic separation of lipids. The separation was achieved using a Waters Acquity UPLC CSH C18 column (2.1 × 150mm with particle size of 1.7μm, part no. 186005298), operating at 55 °C with a flow rate of 200μL/min. The LC gradient consists of a binary buffer system, namely buffer “A” (LC/MS grade water:ACN, 40:60 % V/V) and buffer “B” (IPA:ACN, 90:10 % V/V) both containing 10mM ammonium formate. Independent buffers systems were used for positive and negative ESI modes respectively, for ESI+ the pH of buffers was adjusted using 0.1% formic acid and for negative using 0.1% ammonia solution. The LC gradient was the same for both polarities, namely 60% “B” at T0 hold for 1.5 min, linear increase to 85% “B” at 7 min, increase to 95% “B” at 12.5 min and hold for 4.5 min before returning to starting conditions and holding for further 4.5 min (column stabilization). The voltage applied for ESI+ and ESI- was 3.5kV and 2.5 kV respectively. Injection volumes used were 3 μL for ESI+ and 5 μL for ESI-.

The HESI conditions for 200 μL were as follows: sheath gas 35, auxiliary gas 7 and sweep gas of 0. Ion Transfer tube temperature was set at 300°C and vaporizer temperature at 275°C. These HESI conditions were applied to both metabolomics and lipidomics and lipidomics assays.

#### 2.4.3 Mass spectrometry acquisition

Mass spectrometry (MS) data were acquired using the AcquieX acquisition workflow (data dependent analysis). The MS operating parameters were as follows: MS1 mass resolution 60K, for MS2 30K, stepped energy (HCD) 20, 25, 50, scan range 100-1000, RF len (%) 35, AGC gain, intensity threshold 2e4, 25% custom injection mode with an injection time of 54 ms. An extraction blank was used to create a background exclusion list and a pooled QC was used to create the inclusion list.

#### 2.4.4 Data processing

The metabolomic positive and negative data sets were processed via Compound Discoverer™ (version 3.2) using the untargeted metabolomic workflow with precursor mass tolerance 10 ppm, maximum shift 0.3 min, alignment model adaptive curve, minimum intensity 1^6^, S/N threshold 3, compound consolidation, mass tolerance 10 ppm, RT tolerance 0.3 min. Database matching were performed at MS2 level using Thermo Scientific mzCloud mass spectral database with a similarity index of 50% or higher.

The lipidomic positive and negative data sets were processed via Thermo Scientific LipidSearch™ (version 4) using the following workflow: HCD (high energy collision database), retention time 0.1 min, parent ion mass tolerance 5 ppm, product ion mass tolerance 10 ppm. Alignment method (max), top rank off, minimum m-score 5.0, all isomer peaks, ID quality filter A and B only. Lipid IDs were matched using LipidSearch™ in silico library at MS2 level. Corresponding metabolomics and lipidomics pooled QCs samples were used to assess for instrumental drifts; the relative standard deviation (RSD) variation across the QCs for metabolomics and lipidomics were less than 15%. Any metabolite/lipid feature with an RSD of 25% or less within the QCs was retained.

### 2.5 Proteomics

Proteomics results from a pilot study conducted on the same samples used in this study were previously published and discussed in Mickleburgh et al.^11^. Analyses were conducted following an adapted protocol developed by Procopio and Buckley^39^ for protein extraction and LC-MSMS analysis. MS data for proteomic analysis were made available via ProteomeXchange Consortium via the PRIDE^40^ partner repository with the data set identifier PXD019693 and 10.6019/PXD019693.

### 2.6 Statistical analysis

Metabolomics and lipidomics data were normalised by mean values, cube transformed, and Pareto scaling was applied. Proteomics data were normalised using log2 transformation. The evaluation of the effect of deposition contexts on the omic profiles was carried out using Data Integration Analysis for Biomarker discovery using Latent variable approaches for Omics studies (DIABLO)^41^ based on multiblock sparse PLSDA using the ‘mixOmics’ package in R (version 4.1.2)^42^. The same package was used to evaluate the effects that PMI has on the multiple-omics data. The initial model was tuned using a 3-fold/100 repeats cross-validation to perform parameter selection and produce a final model that maintains the maximum covariance reducing the number of the compounds used for the classification. Enrichment analysis was carried out considering pre- and post-placement samples as well.

## 3 Results

A total of 24 human bone samples were included in this study via LC-MS/MS to obtain a full metabolite, lipid, and protein profile of each sample. Following preliminary inspection via PCA, lipidomics ESI+ results were excluded due to their poor contribution to the discriminant model (**Fig. S1**). The metabolite matrices resulting from the combination of metabolomics ESI+ and ESI-data, resulting in a final matrix with a total of 104 identified compounds after the removal of non-endogenous metabolites as previously described. Non-endogenous compounds were queried in HMDB and excluded from the model. Additionally, 170 lipids, profiled in ESI-, and 132 proteins were considered for further statistical processing. Feature selection using the DIABLO method aims to identify highly correlated and/or discriminant variables across the three omics. Arrow plot (**Fig. 2A**) show the overall separation between fresh and skeletonised samples, which is mainly developed along the first component. However, it is possible to note that the individual with the longest PMI (D4, 872 days) also clusters away from the remaining skeletonised samples along the second component (**Fig. 2B**). Considering the individual -omics consensus plots in **Figure S3**, metabolite and lipid blocks show a better segregation between the individuals D1, D2 and, D3 in their skeletonised state.

**Figure 1.**
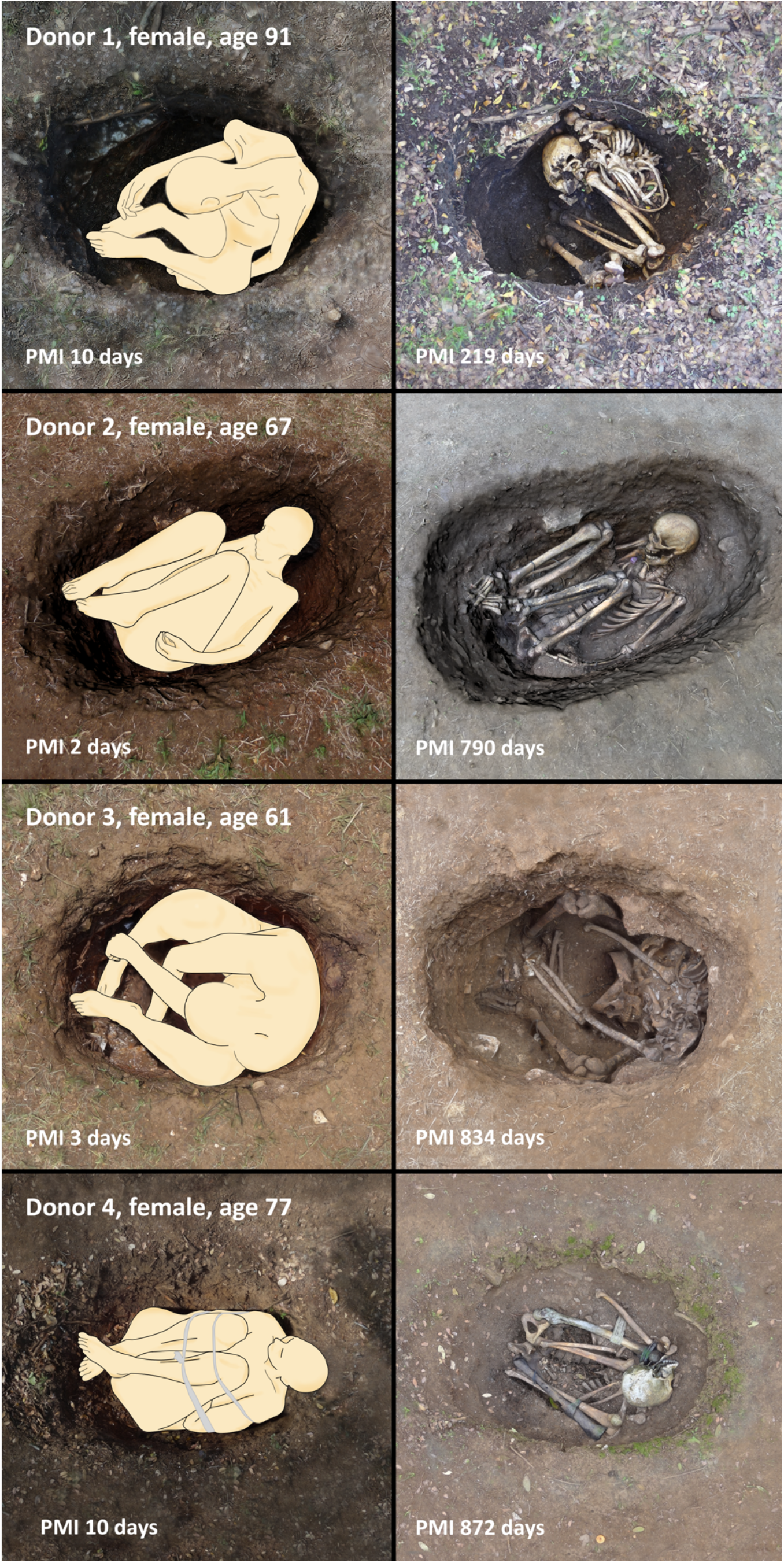
Positioning of the bodies in the single graves (left) pre-decomposition and (right) after complete skeletonization.

**Figure 2.**
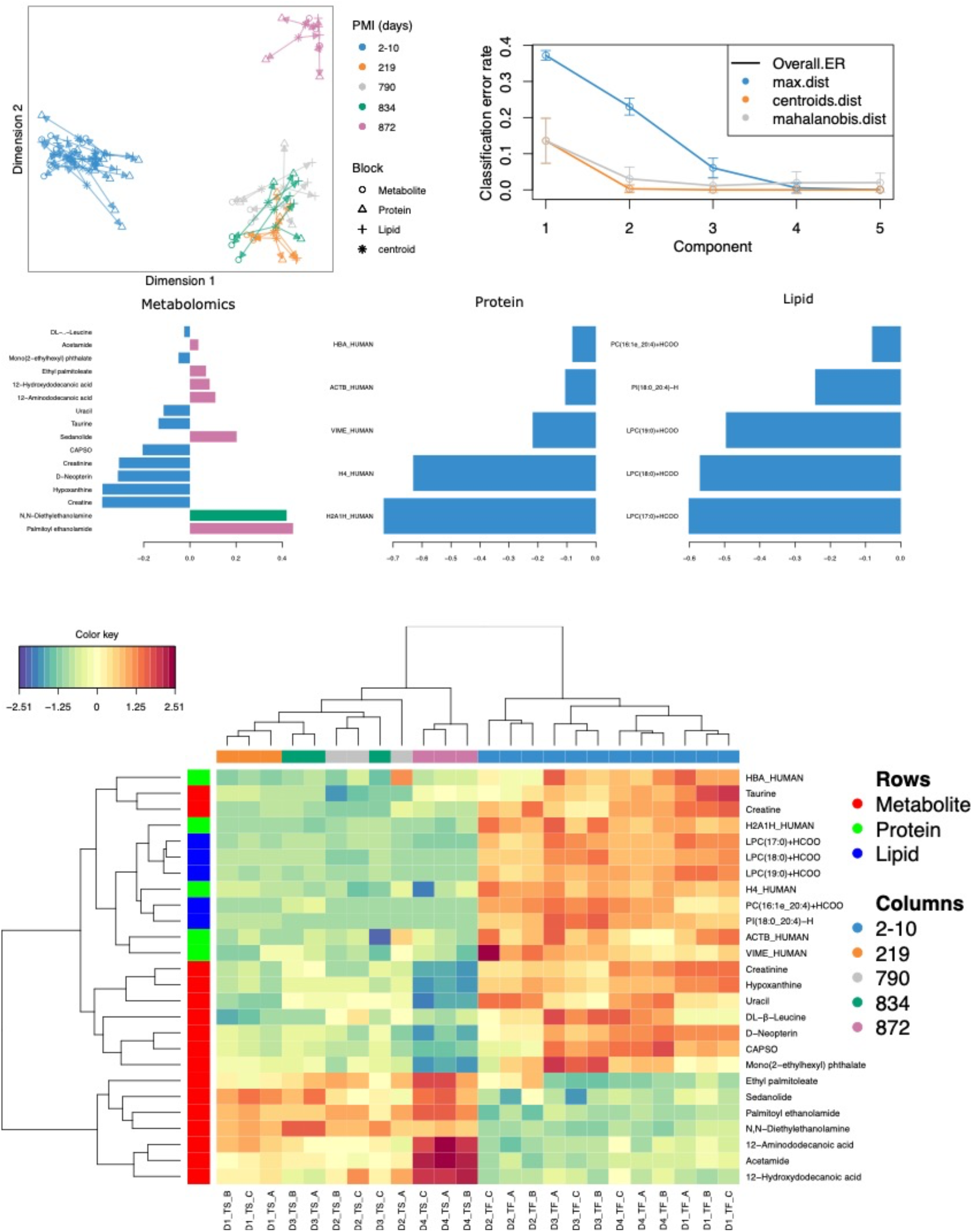
Results for the tuned model. (A) Arrow plot showing multiblock contexts for the overall model. (B) optimal number of components to explain model variable calculated via cross-validation. (C) Loading plot showing how each variable contribute to the covariance of each group. (D) The CIM shows the selected compounds in the final model. It is possible to see that most compounds decrease in intensity after decomposition except for few metabolites and two lipids that specifically increase in certain PMI intervals

The optimal number of components was set at three by means of 3-fold cross-validation repeated 100 times (**Fig. 2B**). The overall balanced error remains below 4% (**Fig. S2**). After tuning the model by attributing the same weight to all the omics blocks, the ideal panel of markers selected in the first component that retains most of the covariance of the system includes 16 metabolites, five lipids and five proteins (**Fig. 2C**). These loading plots show that a few metabolite markers have a high loading for different PMIs, whereas both lipid and protein markers have high values particularly for the fresh samples.

Multi-omics sample variations between bones from fresh and skeletonised cadavers are also supported by the clustered image map (**Fig. 2D**), which shows a clear separation between the two groups. Most of the compounds selected by the model are highly abundant in the fresh samples and less abundant in the skeletonised ones, although the lower panel of metabolites (in **Fig. 2D**) shows an opposite trend. In general, it can be observed that the samples with shorter PMIs (up to 834 days) show a decline in all the proteins, lipids, and for nine of the metabolites selected for the PMI model and an increase in the remaining seven metabolites in comparison with their fresh counterparts. Whereas the decline in the abundance of proteins and lipids in comparison with the fresh samples is similar between all the 12 skeletonised samples, the increase or decrease in the abundance of specific metabolites is more exacerbated in the samples with the longest PMI (872 days) in comparison with the others (**Fig. 2D**).

Evaluating single markers, it is possible to identify those metabolites that increase or decrease consistently across the PMI (**Fig. 3**). More specifically, palmitoyl ethanolamide, N,N-diethylethanolamine, sedanolide, 12-aminododecanoic acid and acetamide show the lowest values for the fresh samples and increasing values with prolonged decomposition time. Another peculiar behaviour is the one shown by the amino acids: taurine, DL’β−leucine, and L−phenylalanine, which drop after the first PMI timepoint, and then moderately increase again. The remaining polar metabolites decrease consistently with PMI with a considerable drop between the baseline and 219 days. Lipids and proteins selected for the model, instead, are all characterised by a drastic reduction in their intensity in the skeletonised samples in comparison with the fresh ones. Proteins selected here are two histone proteins (histone H2A type 1-H (H2A1H), and histone H4 (H4)), haemoglobin subunit alpha (HBA), vimentin (VIME), and actin (ACTB).

**Figure 3.**
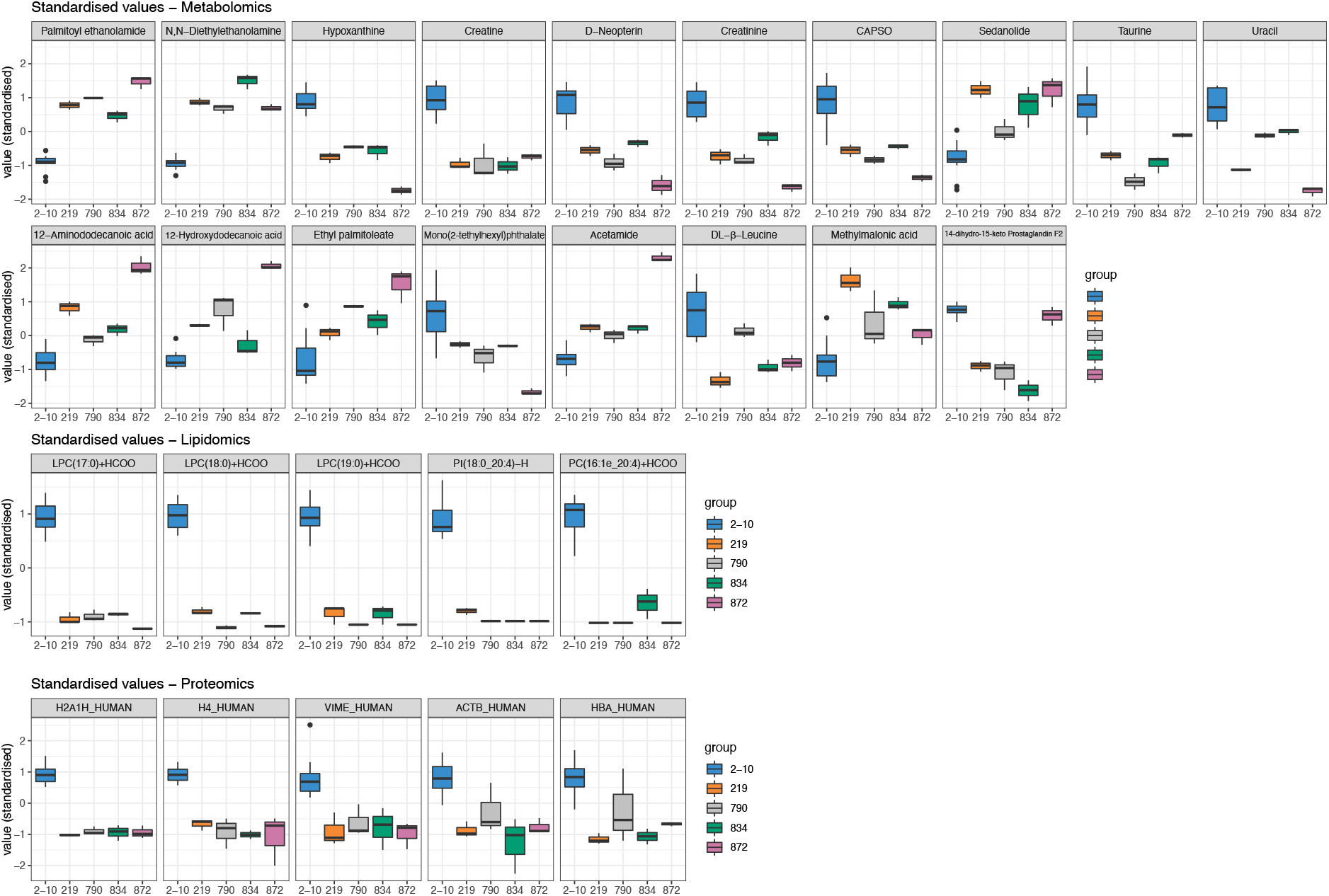
Boxplots of the selected variables after tuning that shows variation with PMI. Variables are expressed in standardised values.

High significant correlations (r>0.9) were also identified between compounds belonging to the three distinct omics blocks (**Fig. 4**). In particular, palmitoyl ethanolamide is negatively correlated with two histone proteins H2A1H and H4 histone proteins, as well as with LPC(17:0)+HCOO, LPC(18:0)+HCOO, LPC(19:0)+HCOO and PI(18:0_24:4)-H. The same metabolite also shows negative correlations with hypoxanthine, creatinine and neopterin. Creatine is positively correlated with LPC(17:0)+HCOO, LPC(18:0)+HCOO and LPC(19:0)+HCOO. D-Neopterin is positively correlated with both histone proteins, with LPC(17:0)+HCOO, LPC(18:0)+HCOO, LPC(19:0)+HCOO and PI(18:0_24:4)-H and with hypoxanthine. Finally, hypoxanthine is positively correlated with LPC(17:0)+HCOO, LPC(18:0)+HCOO, LPC(19:0)+HCOO and with the two histone proteins. Additionally, the five lipids showed positive correlations with each other as well as with the two histone proteins.

**Figure 4.**
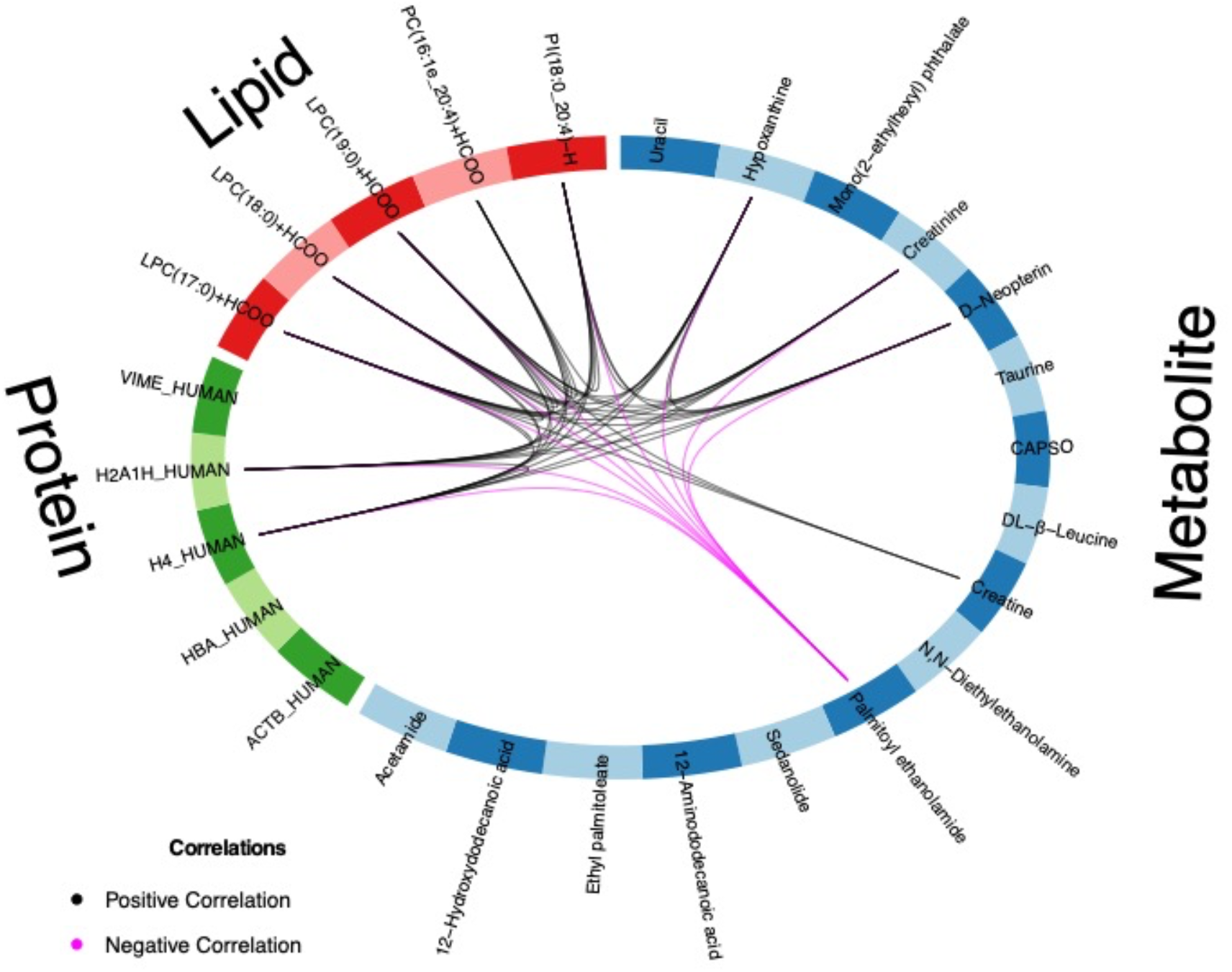
Correlation between different omics blocks highlighting the correlations (at r=0.90) between different compounds obtained with the three omics selected in the final discriminant analysis model.

**Figure 5.**
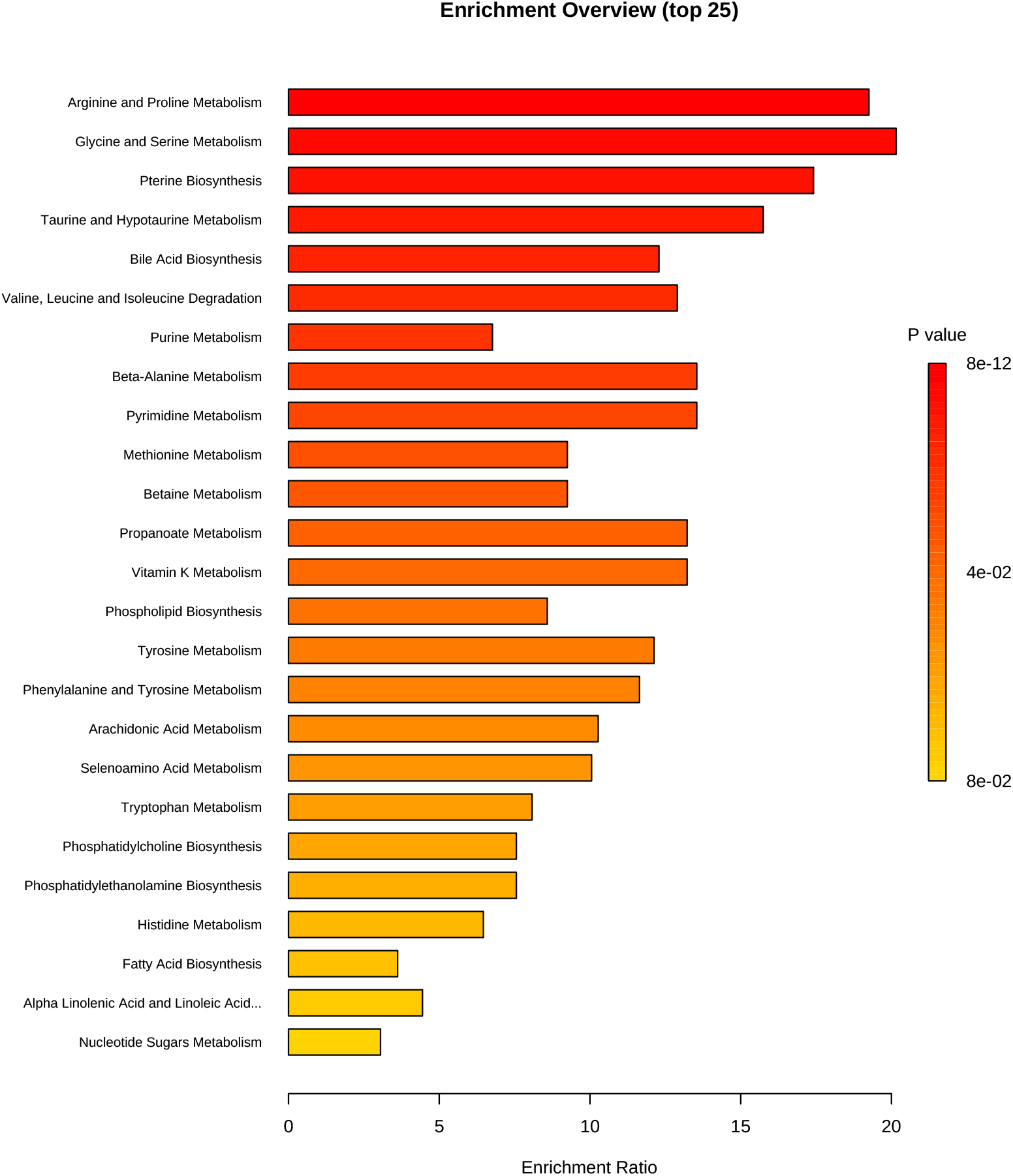
Metabolite set enrichment analysis based on differentially expressed metabolites identified in bone.

## 4 Discussion

This study comprises, to the best of our knowledge, the first attempt to apply a panel of multiple different omics methods to human bone from a controlled decomposition experiment, to identify potential biomarkers for biomolecular post-mortem interval (PMI) estimation. To develop and validate multi-omics PMI estimation methods for forensic application, replication studies in substantial sample sizes of human bone will be necessary. However, the availability of bone samples both before and after decomposition from the same individuals is currently very limited. The work presented here represents a proof-of-concept study on the potential advantages of combining different omics for PMI estimation. The small number of individuals included is consistent with numbers generally used in human decomposition experiments, in which for practical and ethical reasons larger samples, such as used in clinical studies, are very difficult to obtain. While the sample size used here is not suitable for validation purposes, it serves to demonstrate the value and potential of the “ForensOMICS” approach.

As can be seen in **Figure S1**, except for the lipidomics assay acquired in ESI+, the remaining blocks show a very similar pattern in the individual plot. However, on closer inspection, the proteomic profile appears to show quite a considerable overlap between the individuals from three post-decomposition groups (i.e., 219, 790 and 834 days) suggesting that this does not provide sufficient sensitivity to be able to segregate close PMIs. This could be due to the nature of these biomolecules; proteins, in fact, have been seen to be highly stable and could be employed for long-term PMI estimation in forensic scenarios^12,13^ and even in archaeological investigation of skeletal remains^43,44^. Therefore, it possible that the close and rather short PMIs included in the present study could be not fully differentiated using proteomics. Employing the system biology approach for PMI estimation for forensic purposes, by combining more than one class of biomolecules that have different post-mortem stability^17^, provides a biological explanation of the process under investigation. This is achieved in this study by combining different layers of omics (i.e., metabolomics, lipidomics and proteomics) to reconstruct the molecular profile of the system. The DIABLO model simultaneously identifies important markers to optimise classification of PMIs and combines multiple omics techniques^41^. This is normally used to explain the biological mechanisms that determine a disease and its development, while in our case the main advantage is represented by the potential of selecting a pool of compounds that effectively explains, and could accurately estimate, PMI changes over an extended period of time. As an example, most of the studies involving metabolomics for PMI estimation involve quickly degradable matrices (e.g., muscle, blood, humour) collected over a short period of time (<1 month)^14,15,34,45,46^. As previously mentioned, the analysis of proteins in bone have shown applicability to forensic PMI estimation^12,29,47^ as well as to answer archaeological question^48–52^, suggesting the longer survival of this type of biomolecules. Finally, according to the studies presented so far, it seems that post-mortem changes of lipids could provide PMI estimation across several years, although there is great need for validation^27,53^. The combination of these biomolecules’ classes in a multi-omics model could therefore be beneficial for estimating PMI across a broader range of potential PMIs. Metabolites and lipids offer accuracy in the short to medium term while proteins could be the main markers for longer PMIs due to their greater stability. Furthermore, variable selection^41,42^ would offer the advantage of simplifying experimental procedures and targets those markers that behave consistently with PMI. To limit the potential effects of interindividual variability, we considered variables that showed no outliers among the four body donors and created a model that limits as much as possible the number of predictors without affecting the assessment of the PMI.

Our results for the metabolomics assay display clear differences between the pre- and post-placement bone metabolomic profiles, suggesting the potential to use these profiles to assess long PMIs. The small sample size in this study does not allow us to make any deep inferences about the biological significance of the metabolomics profiles of the post-placement samples, as these may have been influenced by exogenous factors. With regards to the pre-placement samples, the PMIs ranging between 2-10 days at 4°C would have allowed some minimal post-mortem modifications in the metabolome to occur^21^. The metabolomic profiles of these samples are characterised by creatine, taurine, hypoxanthine, 3-hydroxybutyrate, creatinine, and phenylaniline. Hypoxanthine is a well-known hallmark of ATP consumption and, consequently, a sign of exhaustion of normal substrates (*i*.*e*., glucose and pyruvate) of the Tri-Carboxylic Acid (TCA) cycle. In conjunction with the presence of creatine, taurine, creatinine, phenylalanine, and 3-hydroxybutyrate, we may hypothesise a switch towards TCA cycle anaplerosis through aminoacidic and ketonic substrates, in pursuit of a resilient ATP production during the early/mid PMIs. Not only was the proposed metabolomic approach able to identify the pre- and post-deposition groups according to the bone metabolome modifications, but it was also sensitive enough to detect at very long PMIs. The presence of exogenous compounds (i.e., caffeine, ecgonine, dextromethorphan, tramadol N-oxide, penbutolol, salicylic acid) that could reflect lifestyle habits or pharmacological therapies, and thus potentially has major implications in forensic toxicology and personal identification, is consistent with evidence from animal models^22^.

Several polar metabolites identified in this study have previously been found in other tissues to show a consistent decay pattern after death. In fact, most of the compounds of interest matched here have already been flagged in other tissues as good potential biomarkers of PMI across shorter timeframes (**Fig. 4**). Uracil, a pyrimidine base of RNA, was previously seen to increase over a 14-day PMI in human muscle tissue when analysed by LC-MS^14^. Similar results for this compound were found in GC-MS analysis of rat’s blood^54^. In contrast, no clear association between this metabolite and PMI was found in aqueous humour^15^. In the present study, after a drop in normalised intestines between the baseline and first PMI, we detected an increase until 834 days, and a drop towards the longest PMI considered. It is worth mentioning that most metabolites drop significantly after the baseline (“fresh”) times (**Fig. 3**), suggesting that compound decomposition is driving this first part of the PMI following the stop of human metabolism. It is interesting that with the increase in PMI there is also an increment in several compounds that could be associated with the breakdown of larger biomolecules (e.g., proteins) or with the presence of microbial communities that leave their own metabolic profile on bone surface. Another common marker of interest is hypoxanthine for its association with hypoxia^15,16,55,56^, that seems to drastically drop between the baseline times and the first PMI timepoint, as well as in the last time interval, showing a good consistency with PMI. In contrast, hypoxanthine was seen to increase until 48 hours and then to decrease at 72 hours in rat blood^57^. Zelentsova et al.^16^ showed a positive relation between hypoxanthine and PMI in human serum, aqueous and vitreous humour. To fully understand the behaviour of this compound in bone tissue, a longitudinal study should be performed also including short PMIs. Leucine has also been reported in short time scale to increase in human muscle tissue^14^ and this agrees with our results where, after the initial drop, we noticed a consistent increase from the first PMI onwards. What can be clearly seen in **Figure 3** is that D2 affects the linearity of the trend, suggesting that there might be some degree of interindividual variability. This is the case for several compounds; this limitation could be mitigated by increasing the number of individuals per timepoint in future studies. Creatinine has previously been reported to be a good marker in both muscle tissue^14^. Although it has not been mentioned in literature previously, we also found that neopterin, a biomarker for immune system activation commonly profiled in blood, serum, and urine^58,59^, has a strong negative correlation with PMI. Taurine, also in accordance with studies on vitreous humour^15^, showed a predictable positive behaviour with PMI. Acetamide is a nitrogen-based compound associated with active and advanced decay^60^ that, not surprisingly, showed the best positive association with PMI, resulting in being the most reliable biomarker within the entire panel considered.

Palmitoylethanolamide is a carboximidic acid that was shown to accumulate in relation with cellular stress in pig brains post-mortem^61^. These findings agree with our study, which revealed a clear increase of this metabolite with increasing PMIs. N,N−diethylethanolamine, belonging to the class of organic compounds known as 1,2-aminoalcohols, has not yet been highlighted for its potential in PMI estimation. In the current study, there is a clear increase of this molecule in the decomposed samples, although no clear trends were observed across the various PMIs. A proposed mechanism for its accumulation is the partial oxidation driven by bacterial decomposition of monosaccharides into organic alcohols^17,18^.

12−aminododecanoic acid and 12-hydroxydodecanoic acid are instead medium-chain fatty acids that show a positive relationship with PMI. Previous studies based on skeletal muscle tissue reported a decline in very-long-chain fatty acids^25,26^ in very short PMIs. It is not possible to exclude that the cleavage of longer chains by the action of lipases or microorganic activity^17,19^. The last compound selected in the final model is methylmalonic acid, a carboxylic acid which is an intermediate in the metabolism of fat and proteins. It has been shown that abnormally high levels of organic acids in blood (organic acidaemia), urine (organic aciduria), brain, and other tissues lead to general metabolic acidosis^62^. In this study, even with a post-mortem increase in its concentration, it is not possible to identify a clear trend across the decomposed samples; this may be related to inter-individual biological differences of the donors involved in this study (e.g., age and health condition).

From the lipidomic assay, only five markers were selected in the final model. These are three lysophosphatidylcholines (LPCs), one phosphatidylcholine (PC) and one phosphatidylinositol (PI), all showing decreasing intensities in the decomposed samples in comparison with the “fresh” ones. PCs are generally the most abundant neutral phospholipids and represent the main constituent in cellular membranes. LPCs are derived from the hydrolysis of dietary and biliary phosphatidylcholines and are absorbed as such in the intestines, but they become re-esterified before being exported in the lymph^63^. They are present in cell membranes and in blood. Their half-life in vivo is limited because of the quick metabolic reaction that involves lysophospholipases and LPC-acyltransferases^64^. In contrast, PLS are amphiphilic molecules that are also minorly present in cell membranes, whose role is to modulate the membrane curvature and to have other bioactive functions such as interacting with peripheral proteins^65^ and inhibiting osteoclast formation^66^. After death, these compounds can be converted into fatty acids via hydrolysis to then hydrogenise or oxidase to form saturated and unsaturated fatty acids^17^. This process is driven by intrinsic tissues lipases^17^. A very limited number of studies have applied lipidomics for PMI estimation. Langley et al.^25^ evaluated human skeletal muscle tissue from 31 donors over a PMI of 2,000 accumulated degree days showing consistent extraction of phosphatidylglycerol (PG) 34:0 and phosphatidylethanolamine (PtdE) 36:4, which showed good correlation with PMI. Wood and Shirley^26^ investigated the lipidome of human anterior quadriceps muscle from one donor at 1-, 9-, and 24-day PMIs showing the decline of sterol sulphates, choline plasmalogens, ethanolamine plasmalogens, and phosphatidylglycerols and the increase of free fatty acids. Our results lend support to these earlier findings and further confirm the potential of lipidomics for PMI estimation. Nonetheless, direct comparison with these studies is not possible as they considered different tissues for much shorter PMIs. Additionally, lipids profiled from the muscle tissue after decomposition are suggested to derive from cell membrane breakdown^25,26^. We suggest that, in bone material, the lipidome under investigation accounts not only for cell membrane decomposition of embedded osteocytes but also for the marrow and fluids embedded in the bone pores. The proteomics results revealed that two ubiquitous proteins (histones), haemoglobin, actin and vimentin are the best candidates within this multi-omics PMI model. These five proteins selected by the model represent those which were best able to discriminate between the “fresh” bones and the “skeletonised” bones but are therefore not necessarily the best biomarkers to differentiate between the four post-decomposition PMIs. For insights on the most suitable protein biomarkers for differentiating between the longer PMIs, identified by excluding the “fresh” samples, see Mickleburgh et al.^11^ It is not surprising to see that the proteins highlighted in the model are either ubiquitous proteins or blood or muscle tissue proteins, as their abundance would naturally be higher in “fresh” bone than in “skeletonised” bones. The haemoglobin subunit alpha (HBA) is found in red blood cells but is often also identified in bone samples with long PMIs from archaeological contexts^67^, and its consistent time-dependent degradation has been previously highlighted in skeletal remains using several platforms^68,69^. Furthermore, it has already been reported in skeletal tissue from controlled decomposition studies of animals, and already highlighted as a potential biomarker for PMI estimation^12^. Vimentin (VIME) was also previously reported by Procopio et al.^12^ to be associated with PMI. It is a filament protein abundant in muscle tissue, and therefore its association with bone, particularly with the “fresh” samples, is not unexpected. However, we emphasize that this could also be due to interindividual variability, and that further investigation may clarify the usefulness of VIME to estimate PMI. Actin (ACTB), similar to vimentin, is a structural protein that forms cross-linked networks in the cytoplasmatic compartments and that is strongly connected with the presence of muscle tissue residues. A previous study showed the decrease in myosin contents with increasing PMIs, similarly to what we observed here for ACTB. The remaining two proteins are both components of the nucleosomes, in our study were shown to be drastically reduced in bone tissue also at the first the baseline PMI taken into consideration. In sum, these results allowed the identification of five protein biomarkers which make good candidates for estimation of short PMIs (<900 days) (*e*.*g*., considering time points limited to months post-mortem) and not for years after death for which structural and functional proteins in bone have been shown better targets to employ for PMI estimation^11,13^.

Based on the findings of this exploratory study, we argue that the multi-omic method we adopted here shows considerable potential for the future development of an accurate and precise PMI estimation method for human bone. Further research should focus on increasing the sample size, to ultimately validate the method for application in forensic investigation of skeletonized human remains. Beyond the findings discussed at length above, we emphasize that it is of paramount importance to establish which biomolecules identified here are associated with the human metabolism and degradation, and which are produced by the decomposers’ microbial activity. Controlled taphonomic experiments on human decomposition at human taphonomy facilities provide the opportunity to elucidate biomolecular decomposition of human bone. A comprehensive understanding of the origin of different compounds is key to provide a detailed explanation of the post-mortem changes that affect bone and other tissues, ultimately helping to shed a light on biomolecular PMI investigations and on the real potential that multi-omics analyses can have in this direction.

## 5 Conclusions

In conclusion, our results support the potential for developing an accurate and precise multi-omics PMI estimation method for human bone for application in forensic contexts to aid criminal investigation and assist with identification of the deceased. Despite the small sample size used here, this study demonstrates how the approach can discriminate between short- and long PMIs. This method can produce classification models including different markers (e.g., protein, metabolites, and lipids) to assess both short- and long-term PMIs, with a high level of accuracy, as the compounds under investigation have complementary decay rates. The use of different biochemical markers that have different post-mortem stability offers the advantage of covering both short-term PMIs, by including metabolites and lipids, and long-term PMIs, by implementing in the model more stable proteins that consistently degrade after death. This coverage would be impossible to achieve by employing any of these three omics methods alone. Furthermore, this approach provides new insights on the biological processes that occur after death and will help establish whether certain molecules are the result of degradation or of bacterial metabolism, a central question in forensic science. The proposed “ForensOMICS” approach must be validated by analysis of substantial sample sizes in future controlled taphonomic experiments, conducted in multiple different environments as environment represents the main source of variation in human decomposition.

## Acknowledgments

The authors acknowledge the UKRI for supporting this work by a UKRI Future Leaders Fellowship (N.P.) under grant MR/S032878/1, as well as the European Research Council (grant 319209) and the Leiden University Fund (grant 5604/30-4-2015/Byvanck) for supporting the actualistic taphonomic experiment at FARF. We would also like to thank the NUOmics Facility at Northumbria for the mass spectrometry analyses and the donors and their next of kin for allowing the use of donated bodies to perform this research.

## Author contributions

Conceptualization, A.B., H.L.M., D.W. N.P.; methodology, A.B., N.P.; software, A.B.; validation, A.B., N.P.; formal analysis, A.B., N.P.; investigation, A.B.; Data interpretation: A.B., A.C., E.L., N.P; resources, N.P.; data curation, A.B.; writing—original draft preparation, All, writing— review and editing, All; visualization, A.B.; supervision, H.L.M., N.P.; project administration, H.L.M, N.P.; funding acquisition, H.L.M, N.P. All authors have read and agreed to the published version of the manuscript.

## Data availability

This data is available at the NIH Common Fund’s National Metabolomics Data Repository (NMDR) website, the Metabolomics Workbench ^70^, https://www.metabolomicsworkbench.org, with Study ID ST002283. The data can be accessed via Project DOI https://www.metabolomicsworkbench.org/data/DRCCMetadata.php?Mode=Project&ProjectID=PR001463.

The R pipeline has been uploaded as a .txt file in the supplementary information.

## Supplementary information

**Figure S1.**
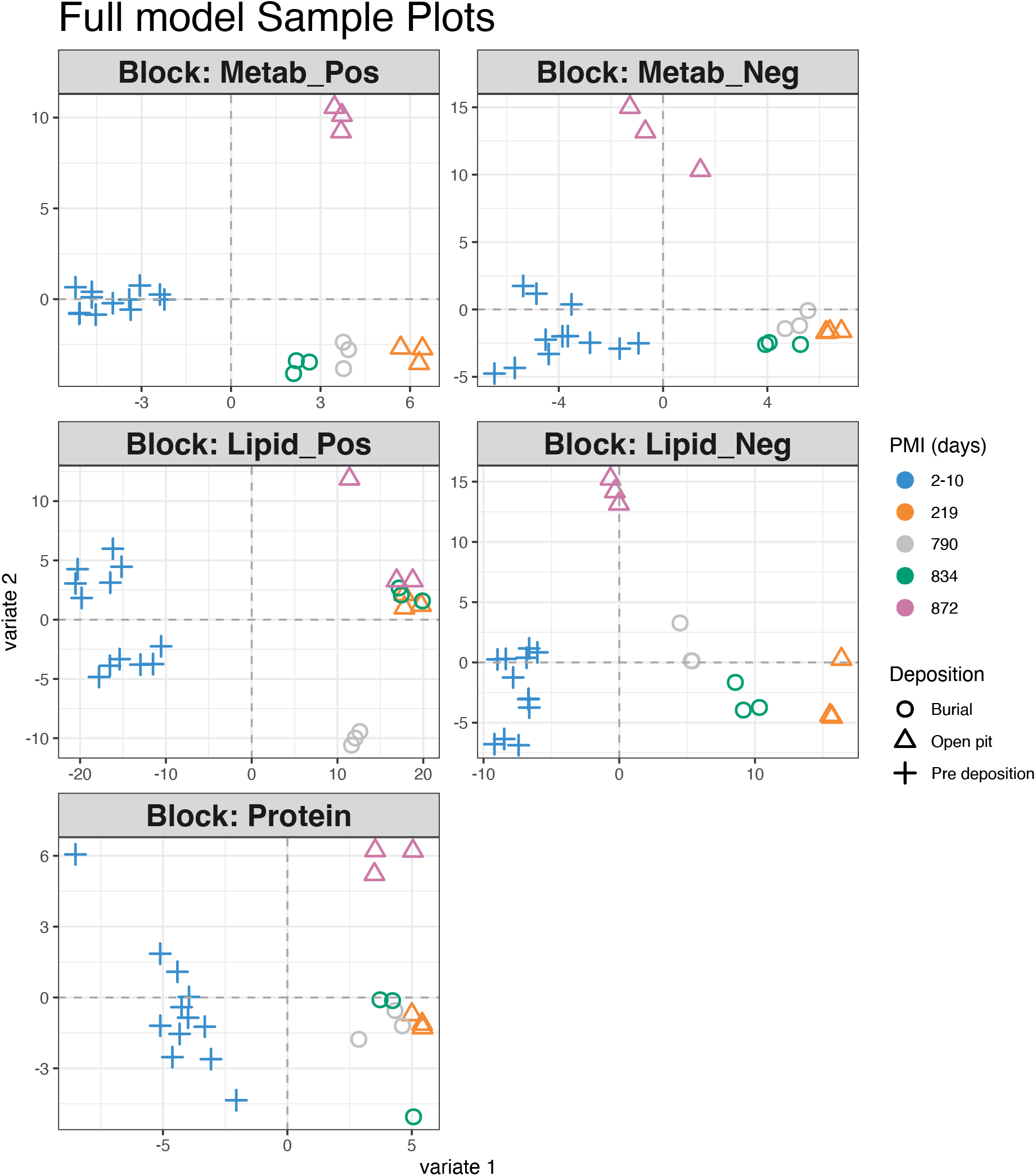
Score plots for PLS-DA results of all the omics blocks considered.

**Figure S2.**
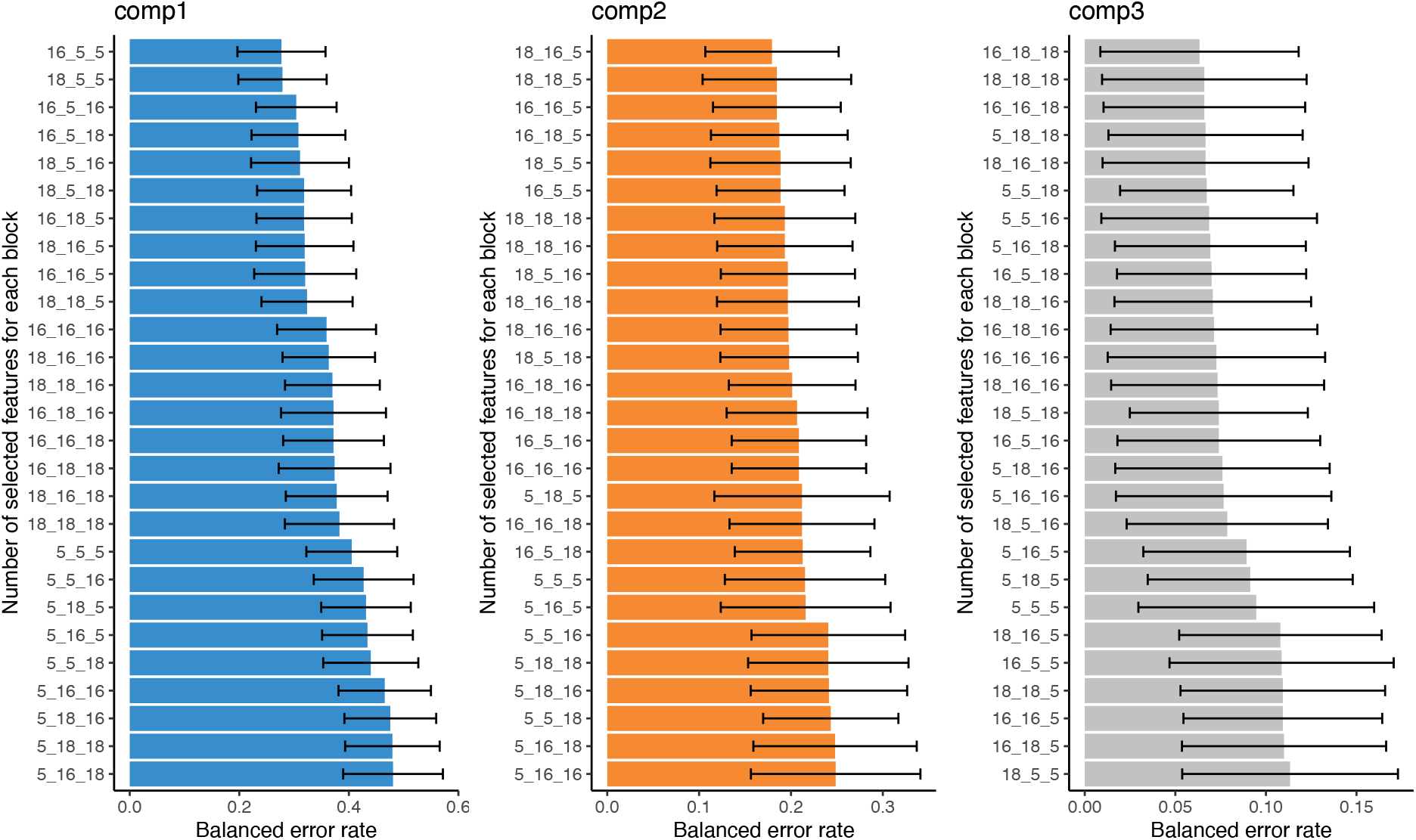
Balanced error variations across variable selection steps.

